# Vasor: Accurate prediction of variant effects for amino acid substitutions in MDR3

**DOI:** 10.1101/2022.02.20.481206

**Authors:** Annika Behrendt, Pegah Golchin, Filip König, Daniel Mulnaes, Amelie Stalke, Carola Dröge, Verena Keitel, Holger Gohlke

**Author notes:** Address correspondence to: Prof. Dr. Holger Gohlke, Institute for Pharmaceutical and Medicinal Chemistry, Heinrich-Heine-Universität Düsseldorf, Universitätsstr. 1, 40225 Düsseldorf, Germany.

## Abstract

**Background / Rationale:** The phosphatidylcholine floppase MDR3 is an essential hepatobiliary transport protein. MDR3 dysfunction is associated with various liver diseases, ranging from severe progressive familial intrahepatic cholestasis to transient forms of intrahepatic cholestasis of pregnancy and familial gallstone disease. Single amino acid substitutions are often found as causative of dysfunction, but identifying the substitution effect in *in vitro* studies is time- and cost-intensive.

**Main results:** We developed Vasor (**V**ariant **as**sessor **o**f MD**R**3), a machine learning-based model to classify novel MDR3 missense variants into the categories benign or pathogenic. Vasor was trained on the, to date, largest dataset specific for MDR3 of benign and pathogenic variants and uses general predictors, namely EVE, EVmutation, PolyPhen-2, I-Mutant2.0, MUpro, MAESTRO, PON-P2, and other variant properties such as half-sphere exposure, PTM site, and secondary structure disruption as input. Vasor consistently outperformed the integrated general predictors and the external prediction tool MutPred2, leading to the current best prediction performance for MDR3 single-site missense variants (on an external test set: F1-score: 0.90, MCC: 0.80). Furthermore, Vasor predictions cover the entire sequence space of MDR3. Vasor is accessible as a webserver at https://cpclab.uni-duesseldorf.de/mdr3_predictor/ for users to rapidly obtain prediction results and a visualization of the substitution site within the MDR3 structure.

**Conclusion:** The MDR3-specific prediction tool Vasor can provide reliable predictions of single site amino acid substitutions, giving users a fast way to assess initially whether a variant is benign or pathogenic.

## 1. Introduction

Bile formation is a carefully regulated system, from bile acid synthesis to secretion of bile acids across the canalicular membrane. Adenosine triphosphate binding cassette (ABC) transporters present on the canalicular membrane of hepatocytes are responsible for the transport of primary bile components, namely bile acids via the bile salt export pump (BSEP, *ABCB11*), cholesterol via the ABC sub-family G members 5 and 8 (ABCG5/ABCG8), as well as phospholipids via the multidrug resistance protein 3 (MDR3). MDR3 encoded by the *ABCB4* gene acts as a floppase, translocating substrates such as phosphatidylcholine from the inner to the outer membrane leaflet (1,2) and exposing the substrate for extraction by bile acids into primary bile (3). Recent studies have suggested different transport pathways, following either an alternating two-site access model through the protein’s inner cavity (4) or a credit-card swipe mechanism along transmembrane helix 7 (5), indicating a need for further research on the exact molecular mechanism. MDR3 dysfunction has been linked to various liver-associated diseases, including intrahepatic cholestasis of pregnancy, low phospholipid-associated cholelithiasis, drug-induced liver injury, progressive familial intrahepatic cholestasis type 3, liver fibrosis/cirrhosis as well as hepatobiliary malignancy (6–12).

It is estimated that at least 70 % of disease-causing *ABCB4* variants are amino acid substitutions, whereas variants leading to premature stop codons and protein truncations are in the minority (13). However, while the advancement of sequencing allows rapid testing of patients, it remains challenging for clinicians and researchers to assess the potential impact of novel missense variants.

Evaluation of newly found MDR3 amino acid substitutions by *in vitro* cellular assays remains time-consuming. Machine learning-based prediction tools instead offer rapid analysis and have led in recent years to a plethora of predictors (14,15). Nonetheless, general predictors do not consistently perform well on all proteins, necessitating the development of protein-specific prediction tools. To date, there is no MDR3-specific predictor available for classifying amino acid substitutions despite MDR3’s vital role in bile homeostasis. An initial evaluation of general predictor performances on MDR3 variants suggested MutPred as a well-performing tool (16); however, generalization is difficult due to only 21 tested variants with established cellular effects. Additionally, the tested variants presented a clear bias towards pathogenic effects.

Here, we created an MDR3-specific variant dataset and trained a machine learning algorithm using several established general prediction tools, namely EVE, EVmutation, PolyPhen-2, I-Mutant2.0, MUpro, MAESTRO, and PON-P2 (17–23), as well as half-sphere exposure, post-translational modification (PTM) site influence, and secondary structure disruption as features to obtain an MDR3-specific prediction tool for help in classifying variants as benign or pathogenic (see Fig. 1 for a graphical overview). Our predictor, Vasor (**V**ariant **as**sessment **o**f MD**R**3), performed better than each integrated general predictor. Additionally, Vasor outperformed MutPred2, a general predictor we chose for comparison based on the suggested high performance of its predecessor MutPred on MDR3 (16). We provide easy access to Vasor via a webserver, where users can enter a missense variant of interest and obtain a prediction if it is benign or pathogenic together with an estimate of the prediction probability. Additionally, the mutation site is displayed on the structure of MDR3, giving the user a comprehensive view of the local site and the overall position of the assessed variant.

**Fig. 1:**
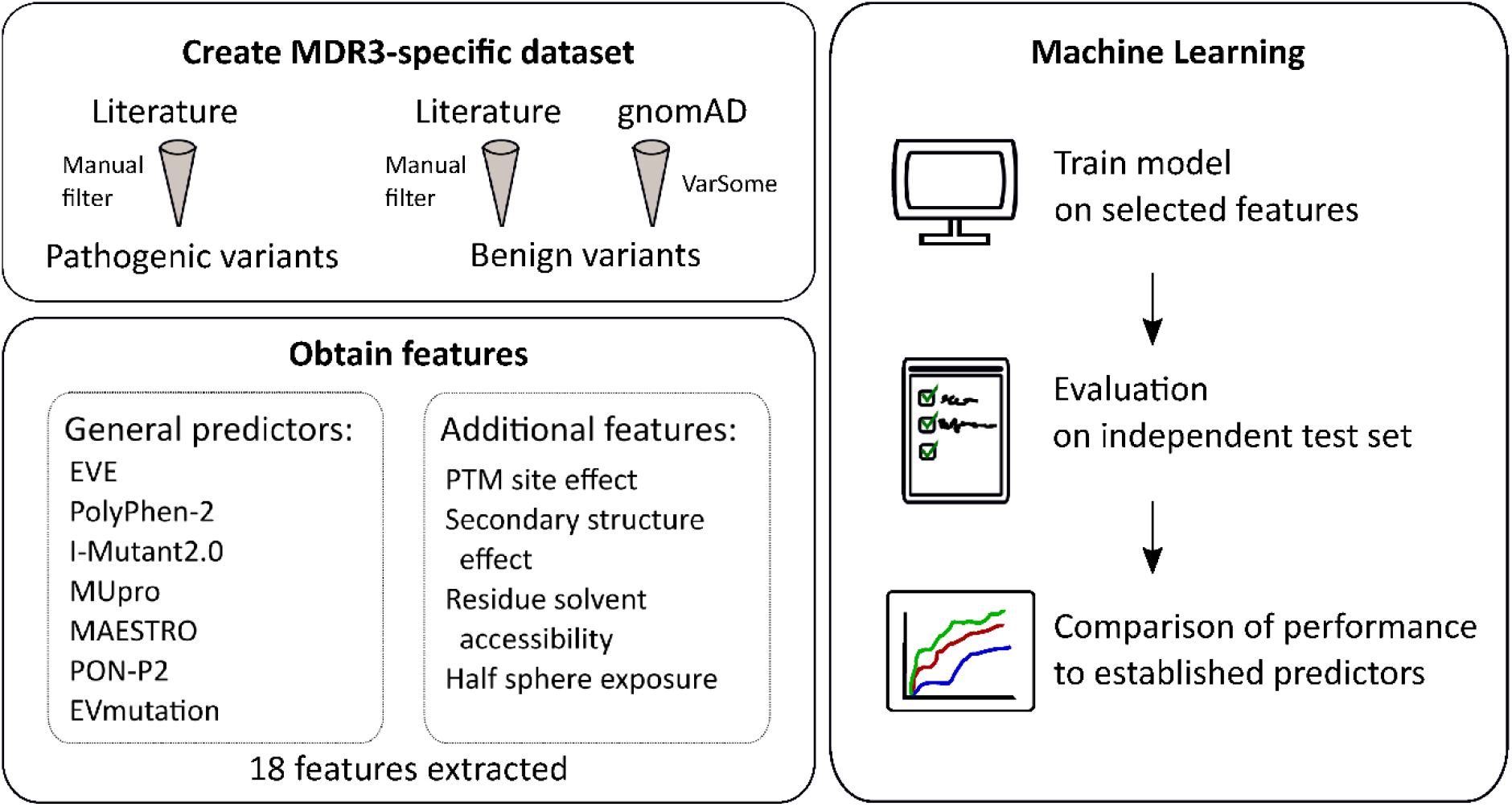
Graphical overview of dataset generation and machine learning approach. For details see text.

## 2. Method and Implementation

### 2.1. MDR3 missense variants

MDR3 variants were obtained from a literature search for variants causative of MDR3 dysfunction or known variants with no effect in any MDR3-associated disease (4,6,10,12,13,16,24–57). We excluded variants with unclear information on disease association (i.e., no *in vitro* verification analysis and no information on clinical indications for disease association) to eliminate False Positives or False Negatives. As studied benign variants for MDR3 are rare (13,16), further missense variants were obtained from gnomAD v2.1.1 (58) to increase the number of benign variants. During the generation of the gnomAD database, individuals with severe pediatric diseases are removed; however, it is possible that pathogenic variants exist in the gnomAD dataset. Accordingly, we employed a selection step to exclude false-negative cases of MDR3 variants. Using the platform VarSome (59), variants were pre-classified following the guidelines of The American College of Medical Genetics and Association for Molecular Pathology (ACMG-AMP) (60) rules, and variants with a likely pathogenic or pathogenic effect were removed, whereas variants with uncertain significance, likely benign, or benign classification by VarSome were integrated into the dataset. These steps were included to create a high-quality dataset to keep the number of misclassified variants low but at the same time retain a sufficiently high number of variants. The final list of variants contained 85 pathogenic and 279 benign variants. Every variant was mapped to the longest MDR3 isoform, corresponding to Uniprot (61) entry P21439-1.

### 2.2. Dataset and features

The list of MDR3 variants was subjected to established general predictors for missense mutations (EVE, PolyPhen-2, I-Mutant2.0, MUpro, MAESTRO, PON-P2, and EVmutation), and additional features (half-sphere exposure, secondary structure, PTM site, and relative solvent accessibility) were computed, creating an MDR3-specific feature set.

EVE (Evolutionary models of Variant Effects) is a recently developed unsupervised computational method, which trained Bayesian variational autoencoders on multiple sequence alignments to classify variant effects based on a variant-specific, computed evolutionary index followed by a fitted global-local mixture of Gaussian Mixture Models (17). PolyPhen-2 employs a Naïve Bayes classifier for predicting variant effects using sequence-based features and structure-based features, if available (18). I-Mutant2.0 predicts protein stability changes, using a Support Vector Machine-based tool trained on either sequence or structural information (19). MUpro predicts stability changes upon single-site mutations, using sequence and structural information if available, using a Support Vector Machine Approach (20). Both I-Mutant2.0 and MUpro predict the direction of stability change and the energy difference. MAESTRO employs a combination of machine learning approaches (neural network, support vector machine, and multiple linear regression) to predict the energy difference introduced by missense mutations based on consensus, along with predicting a confidence score (21). PON-P2 applies selected features from evolutionary conservation and biochemical properties of amino acids to develop a random forest classifier that classifies mutations into benign or pathogenic cases, or those with unknown significance (22). EVmutation explicitly considers interdependencies between residues or nucleotide bases in their unsupervised statistical method to include epistasis (23).

EVE and EVmutation predictions for the MDR3 protein were accessed using the pre-computed dataset available from the creators of EVE (https://evemodel.org/) and Evmutation (https://marks.hms.harvard.edu/evmutation/human_proteins.html), respectively. I-Mutant2.0, Mupro, and MAESTRO predictions were generated using their standalone downloadable versions. PolyPhen-2 predictions were accessed using the batch query of the webserver (http://genetics.bwh.harvard.edu/pph2/bgi.shtml) with the default values. PON-P2 predictions were generated using the sequence submission feature for variants of the webserver (http://structure.bmc.lu.se/PON-P2/).

Additional features were added to explicitly integrate effects on PTM sites, variant location in α-helical or β-sheet secondary structure, and effects on residue solvent accessibility. Known PTM sites from the literature were supplemented by potential PTM sites, which were predicted using PhosphoMotif (62), PhosphoSitePlus (63), NetPhos (61), and the ELM database (65). Secondary structure was extracted from the MDR3 structure (PDB ID 6S7P) using DSSP (66,67). Relative solvent accessibility was computed based on residue exposure calculated with DSSP divided by the maximal residue solvent accessibility calculated by Thien et al. (68). Half-sphere exposure was introduced by Hamelryk (69) to measure residue solvent exposure and surpass limitations of relative solvent accessibility, which does not differentiate between residues closely buried beneath the surface and residues deeply buried. It was implemented using values from the Biopython HSExposure module calculated according to the half-sphere corresponding to the direction of the sidechain of the residue as measured from the Cα atom.

### 2.3. Machine learning

The obtained dataset was cleaned from non-numerical values. In the case of binary features, such as classification features of general predictors, -1 was set if no prediction was available to distinguish from benign (indicated by value 0) or pathogenic (indicated by value 1) predictions. Additionally, relative solvent accessibility and half-sphere exposure were set to -1 if no prediction value was obtained, to distinguish from prediction values of 0. Other numerical features were replaced by 0 if no prediction for the respective feature was available. The correlation between features within the dataset was assessed by the Spearman *R* correlation coefficient.

A test set was generated by selecting 20 benign and 20 pathogenic variants from the overall dataset. To avoid a bias towards specific amino acids, we minimized the root-mean-square deviation (RMSD)-based difference between the amino acid distribution of the variants within the test set compared to the overall dataset (Suppl. Fig. 1): After randomly drawing ten variants into the test set, the RMSD-based difference between the amino acid distribution of the general dataset and current test set was computed; further variants were only transferred into the test set if they met one of the following conditions: (a) the RMSD between reference sequence and substituted amino acid distributions decreased by addition of the new variant, (b) the RMSD between reference sequence amino acid distributions decreased while the RMSD between substituted amino acid distributions did not increase more than 0.1, or (c) the RMSD between substituted amino acid distributions decreased while the RMSD between reference sequence amino acid distributions did not increase more than 0.1. Due to the limited size of the dataset, otherwise, it might not be possible to draw a variant for the test set. The test set was withheld from the machine learning training step and used for final validation.

To handle the imbalance between the pathogenic (85 variants) and benign (279 variants) class, we used the synthetic minority oversampling technique (SMOTE) (70). This method generates new synthetic data points by using existing minority data points within the *N*-dimensional dataset space, drawing lines to the five nearest minority class neighbors, and randomly selecting synthetic data points along these lines to balance out the classes.

On the training dataset, the XGBoost algorithm (71) (as implemented in the python library) was trained using the default gradient-boosted tree (gbtree); the maximum depth of a tree (max_depth) was set to 3, subsample to 0.6, and the step size (learning_rate) to 0.02. The training was evaluated using repeated *k*-fold cross-validation, with *k* set to 3 and the value of repeats (n_repeats) to 5. Using this procedure, the training dataset was randomly split into three equally sized folds, where each fold is used as an internal test dataset with the remaining two folds as training datasets, respectively. The performance results were measured and visualized in receiver-operator curves (ROC) for comparison to the final test set. These steps were repeated five times.

To reduce features and estimate feature importance, we analyzed the tree-based feature importance and the permutation importance, leading to the removal of the four least-informative features shared in both feature importance measures: relative solvent accessibility, I-Mutant2.0 stability sign, I-Mutant2.0 deltaG value, and PON-P2 probability value. Tree-based feature importance was computed using the XGBoost algorithm built-in feature and the ‘gain’ (average gain across all splits where a feature is used). Permutation-based feature importance was computed by random shuffling each feature consecutively, followed by a performance test, denoting performance alterations upon feature permutation. The performance of the model without feature selection is shown in Suppl. Fig. 2.

The trained model, termed Vasor (**V**ariant **as**sessment **o**f MD**R**3), predicts a probability, ranging from 0 to 1, for a given variant to belong to the pathogenic class. Predictions above (below) 0.5 are classified as pathogenic (benign).

### 2.4. Comparison to established predictors

To assess the general performance of Vasor, we compared it to the general predictors EVE, PolyPhen-2, PON-P2, and MutPred2. MutPred2 predictions were used to compare our prediction tool to an ‘external’ general predictor, as MutPred2 was not used as an input feature for Vasor. The standalone version of MutPred2 was used to classify each variant within the entire dataset. The performance of Vasor and the other predictors was evaluated on the entire dataset and the test set. This ensured increased fairness for the performance comparison, as Vasor may have an advantage over other predictors based on its training on the training dataset. ROC and precision-recall-curves were adjusted to the availability of variants each predictor was able to classify over the entire dataset (i.e., if general predictors did not classify a variant into the category “benign” or “pathogenic”, the respective variant could not be assessed, and curves were shown only on assessable variants). To account for this, the coverage of each predictor of the MDR3 dataset was computed.

### 2.5. Performance evaluation

The performance of Vasor and the other prediction tools was evaluated using recommended measures for binary classifiers (72), including additionally the F1-Score as well as visualization in ROC and precision-recall-curves. The measures are based on the values of correctly classified variants, indicated by True Positives (TP) for correctly predicted pathogenic variants and True Negatives (TN) for correctly predicted benign variants, as well as incorrectly classified variants, indicated by False Positives (FP) for variants predicted as pathogenic albeit being benign and False Negatives (FN) for variants predicted as benign albeit being pathogenic. The analyzed measures of recall, specificity, precision, negative predictive value (NPV), accuracy, F1-score, and Matthews correlation coefficient were calculated as follows:

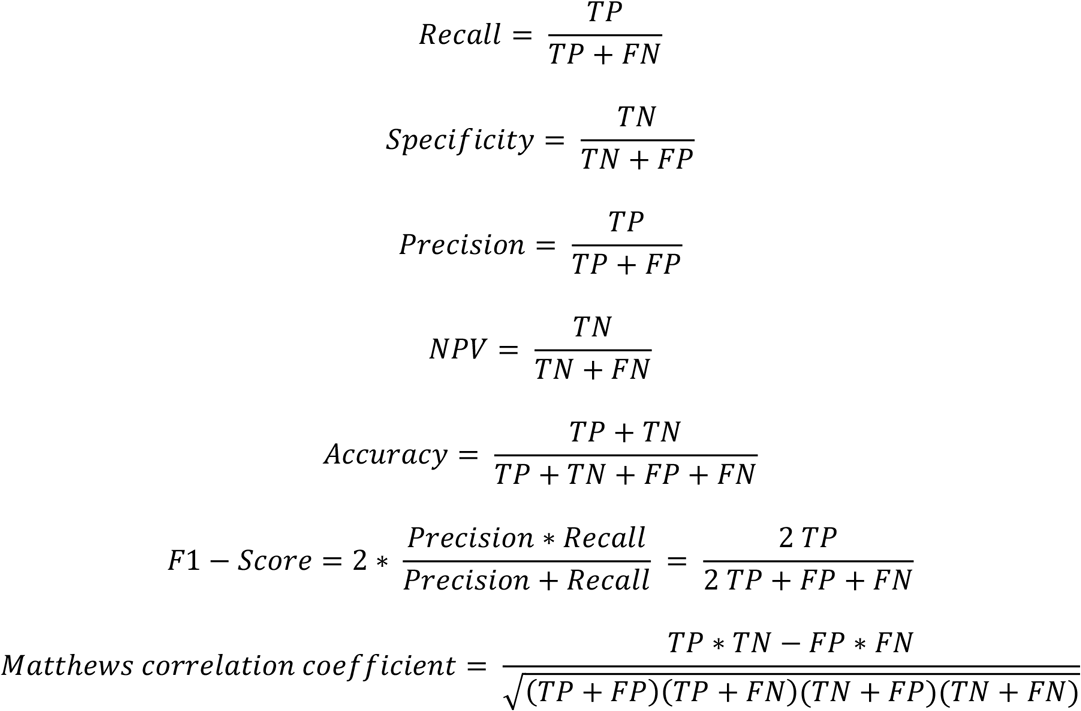

### 2.6. Webserver tool

Vasor can be accessed online via https://cpclab.uni-duesseldorf.de/mdr3_predictor/. Users can enter a single-site amino acid missense MDR3 variant of interest, keeping in mind that the tool will only recognize MDR3 variants corresponding to the largest protein isoform, UniProt ID P21439-1. Further, the entry needs to be in the format of the standard IUPAC code for amino acids, entering first the one-letter code of the amino acid of the reference sequence, followed by the position and the amino acid substitution of interest. On the results page, users can see the predicted classification (either benign or pathogenic) and the probability of the given variant being pathogenic. This probability ranges from 0 (highest probability for the variant to be benign) to 1 (highest probability for the variant to be pathogenic). Probability values close to the cut-off value of 0.5 indicate less confidence in the prediction.

Additionally, the results page displays the structure of the MDR3 protein (PDB ID: 6S7P) with the NGL Viewer (73,74), including the membrane localization obtained from the OPM database (75) as a red and blue plane. The substituted residue is colored according to the predicted effect either in red (pathogenic) or green (benign). The user can download a zip archive containing a high-resolution image of the complete protein, PDB files of the reference sequence and the variant protein, and high-resolution images of the position with the reference sequence residue or the substituted one.

### 2.7. Code availability

The code for Vasor was written in Python 3.9 and will be provided for download at https://cpclab.uni-duesseldorf.de/index.php/Software.

## 3. Results

### 3.1. Generation of a dataset with informative features and good overall coverage of the MDR3 protein

To establish an MDR3-specific prediction tool, we first prepared a dataset of benign and pathogenic MDR3 variants. Relevant literature on MDR3-associated diseases was screened. Variants with unclear association to effects were omitted to avoid misclassified variants within the dataset. In addition, the gnomAD database (58) was screened for MDR3 variants, and the results were subjected to filtering by VarSome (59) using ACMG-AMP rules (60) to remove variants with a high potential for a pathogenic effect. This step was necessary as pathogenic MDR3 variants on a single allele with a potential late-onset or mild phenotype might have been included in the gnomAD database. Next, we used well-established general predictors (EVE (17), EVmutation (23), PolyPhen-2 (18), I-Mutant2.0 (19), MUpro (20), MAESTRO (56) and PON-P2 (22)) and descriptors of the variant site, namely the disruption of secondary structure, possible PTM site disturbance, and changes in the relative solvent accessibility and half-sphere exposure of the position in question, as features in the dataset. Projecting the variant locations from the dataset onto the known cryo-EM structure of MDR3 (PDB ID 6S7P) (4) revealed a broad coverage of the structure with benign and pathogenic variants (Fig. 2A). No functional domain is devoid of variants, and we do not observe large clusters of benign or pathogenic variants, which may be indicative of a potential bias within the dataset. Such a bias might prevent applying the tool to areas of low coverage. Hence, we expect that our tool can generalize predictions to every position of MDR3.

**Fig. 2:**
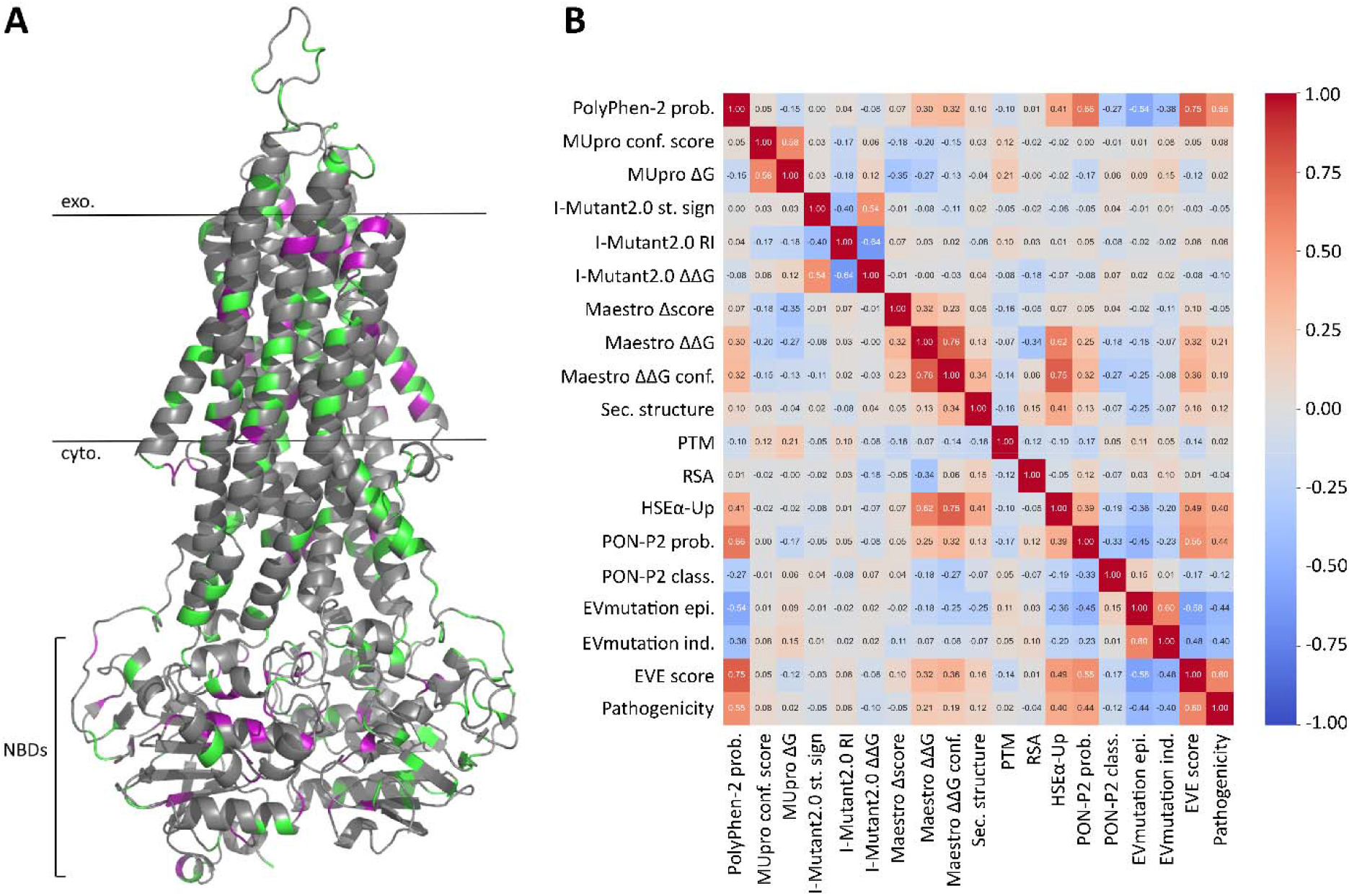
Coverage of MDR3 by the dataset and correlation analysis of features. [A] Mapping of dataset variants onto the MDR3 structure. Benign variants are marked in green and pathogenic variants in magenta. [B] Spearman rank correlation matrix of features computed for the dataset. Abb.: prob. – probability, conf. – confidence, st. sign – stability sign, RI – reliability index, sec. structure – secondary structure, NBD - nucleotide-binding domain, PTM – posttranslational modification, RSA – relative solvent accessibility, HSE – half-sphere exposure, epi. – epistatic, ind. – independent.

To further probe for domains of low applicability, we mapped variants misclassified by Vasor to the MDR3 structure. Misclassified variants from the dataset tend to occur on the solvent-exposed surface of the protein rather than within buried regions of the protein (Suppl. Fig. 3). As solvent-exposed residues are less evolutionary conserved than buried residues (76), the obtained trend might visualize the underlying increased uncertainty of those integrated general predictors that are based on evolutionary sequence conservation. Overall, also given the small number of misclassifications, we do not see indications of domains of increased uncertainty for MDR3 predictions. The correlation coefficients between input features range from -0.64 to 0.76 over the 18 features (Fig. 2 B), indicating that each feature adds information that does not overlap with information from another feature.

### 3.2. Generating Vasor: training the XGBoost algorithm on the dataset

For Machine Learning models to function reliably, it is vital to estimate potential over- or under-fitting of the trained model. One of the most important techniques in that respect is the hold-out method, where a subsection of the entire dataset is split off as an external test set. Ideally, the test set has a similar probability distribution as the entire dataset (77); however, this is not certain if a test set is randomly drawn. Therefore, we paid attention to drawing our test set with a similar distribution of amino acids, as to both reference sequence and variant amino acid distributions, by minimizing the RMSD-based difference in amino acid distributions to the overall dataset. Besides, the test set contained an equal amount of benign and pathogenic variants, 20 each (Suppl. Fig. 1).

Next, for the remaining dataset, SMOTE (70) was used to create synthetic examples of the minority class (pathogenic variants) to balance the classes as binary classifiers otherwise tend to favor the majority class (benign variants) for prediction outcomes. The final training dataset consisted of 259 data points for each class, benign and pathogenic, upon which an XGBoost algorithm was trained. To evaluate the most important features, we measured and visualized feature importance (Suppl. Fig. 4) and removed the four consistently least-important features (Suppl. Fig. 5) without reducing performance. Of note, EVE is highly important for the prediction outcome of the model, indicating that Vasor primarily relies on EVE’s predictions compared to other features.

Performance estimates were visualized within a repeated *k*-fold cross-validation and compared to the performance against the held-out test set (Fig. 3 A). The trained model performs on the test set with an accuracy of 90 %, with 18 out of 20 variants being predicted correctly, both for the benign and the pathogenic class (Fig. 3 B). Notably, the performance based on the *k*-fold cross-validation does not differ from that on the independent test set, indicating a well-fit model without over- or under-fitting.

**Fig. 3:**
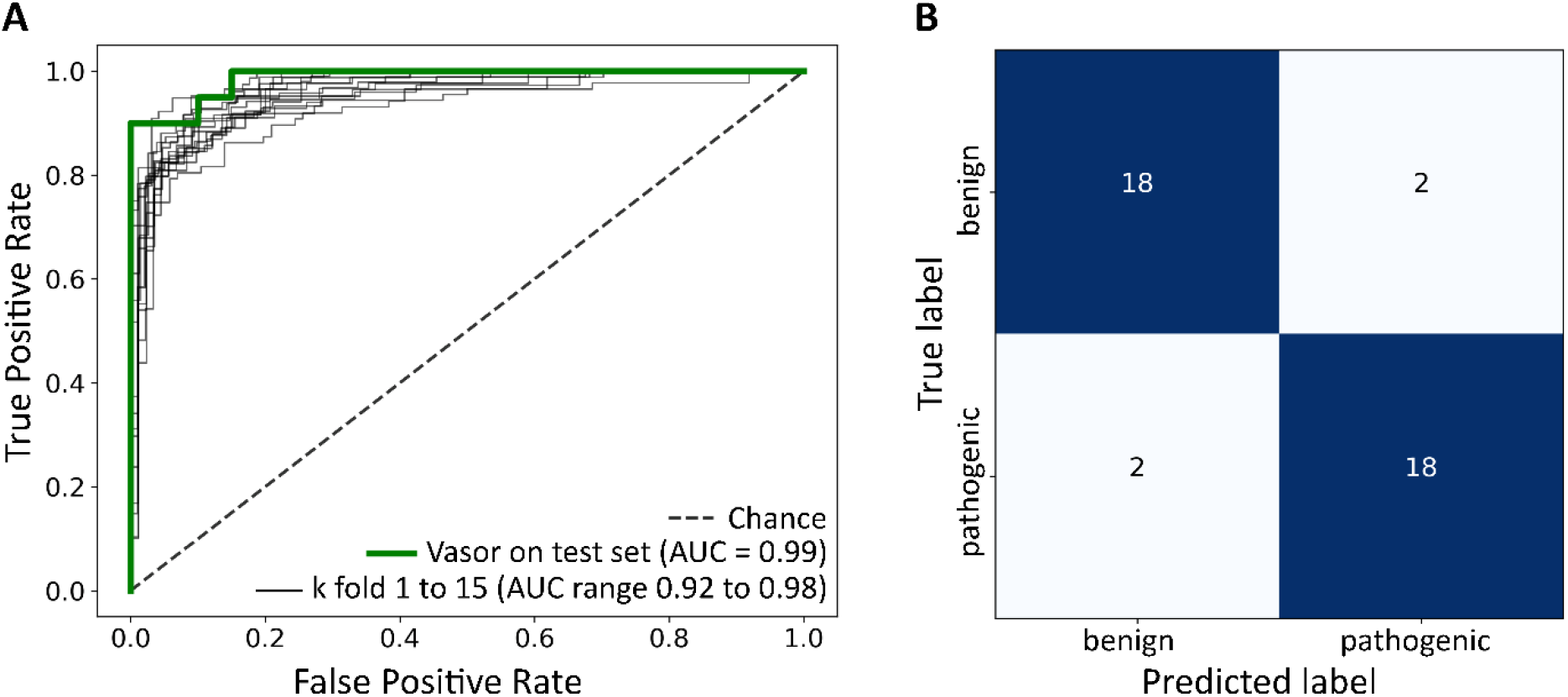
Performance of Vasor on the test set. [A] ROC curve of the Vasor performance on the test set (green line) compared to performance estimates from repeated *k*-fold cross-validation (black lines). [B] Confusion matrix of Vasor performance on the test set.

### 3.3. Vasor outperforms integrated general predictors and the external general predictor MutPred2

We compared the performance of Vasor with general predictors on the entire dataset. We compared Vasor to EVE, PolyPhen-2, and PON-P2, which were integrated as features into the dataset on which Vasor was trained. As such, Vasor should outperform each predictor due to the additional information gathered from the other features. Additionally, we compared Vasor to MutPred2 as an external prediction tool; the predecessor tool MutPred was indicated to perform well on MDR3 classification problems (16). Vasor outperformed EVE, PolyPhen-2, PON-P2, and MutPred2 according to ROC (Fig. 4 A) and precision-recall-curves (Fig. 4 C), with an area under the curve (AUC) of 0.98 for Vasor against 0.90 for EVE, 0.89 for MutPred2, 0.87 for PolyPhen2, and 0.80 for PON-P2 for the ROC and an AUC of 0.94 for Vasor against an AUC of 0.86 for EVE, 0.74 for MutPred2, 0.72 for PolyPhen2, and 0.51 for PON-P2 for the precision-recall-curves. Precision-recall-curves have been shown to be more robust and accurate for binary classifiers on imbalanced datasets (77), like in our case.

**Fig. 4:**
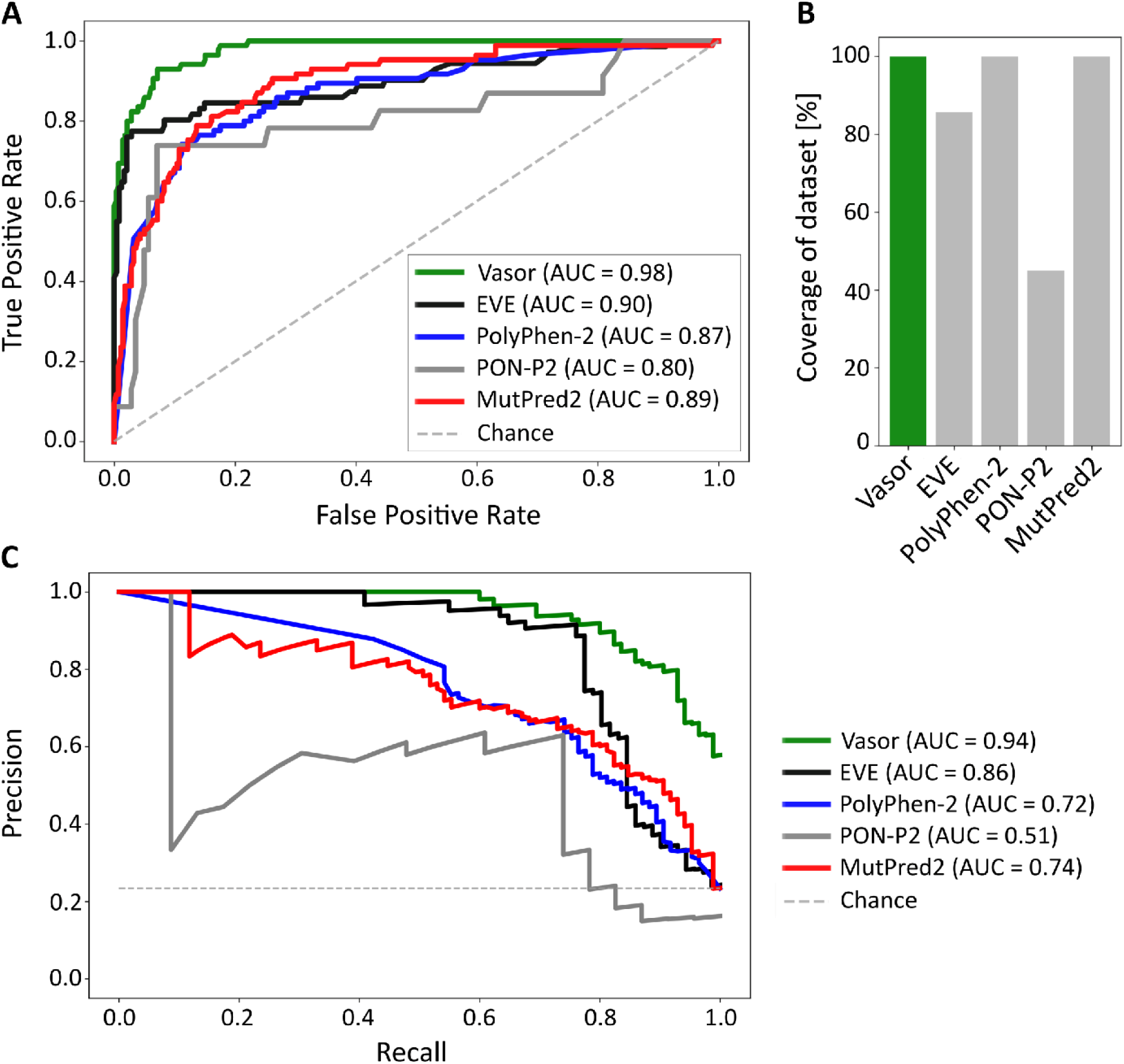
Performance of Vasor in comparison to established general predictors. [A] ROC curve of the performance of Vasor, EVE, PolyPhen-2, PON-P2, and MutPred2 on the variants of the entire dataset. Note that the performance was determined for those variants each predictor was able to make a prediction for (see [B]). [B] Coverage of dataset variants by the predictors. [C] Precision-recall-curves of the predictors. Performance was determined for those variants each predictor was able to make a prediction for.

Noteworthy, the second-best performing predictor, EVE, was the most important feature for Vasor, suggesting that the machine learning model recognized the information contained within this feature as highly correlated with the true output and its value in predicting the output correctly. However, EVE could only predict 85.7 % of the variants in the dataset, whereas Vasor, by design, predicted an outcome for every possible missense variant of MDR3 (Fig. 4 B and Table 1).

**Table 1:** Detailed performance measurements of Vasor in comparison to EVE, PolyPhen-2, PON-P2, and MutPred2 on the entire dataset.

|  | **Vazor** | **EVE** | **PolyPhen-2** | **PON-P2** | **MutPred2** |
| --- | --- | --- | --- | --- | --- |
| **Recall** | 0.84 | 0.73 | 0.84 | 0.74 | 0.52 |
| **Specificity** | 0.96 | 0.98 | 0.74 | 0.89 | 0.95 |
| **Precision** | 0.87 | 0.91 | 0.49 | 0.52 | 0.77 |
| **NPR** | 0.95 | 0.93 | 0.94 | 0.95 | 0.87 |
| **Accuracy** | 0.93 | 0.92 | 0.76 | 0.87 | 0.85 |
| **F1-Score** | 0.85 | 0.81 | 0.62 | 0.61 | 0.62 |
| **MCC** | 0.81 | 0.77 | 0.50 | 0.54 | 0.55 |
| **TP** | 71 | 52 | 71 | 17 | 44 |
| **FN** | 14 | 19 | 14 | 6 | 41 |
| **TN** | 268 | 236 | 206 | 125 | 266 |
| **FP** | 11 | 5 | 73 | 16 | 13 |
| **Coverage [%]** | 100 | 85.7 | 100 | 45.1 | 100 |

Additional performance measures are summarized in Table 1, indicating that Vasor outperforms existing prediction tools according to the weighted measures F1-Score (0.85) and Matthews correlation coefficient (MCC) (0.81). Specifically, Vasor achieved a low number of False Negatives. Comparable low values in False Negatives are achieved by PolyPhen2, but at the cost of an increased number of False Positives, and PON-P2, but only at coverage of 45.1 % of the variants in the MDR3 protein and an increased number of False Positives.

When comparing the performance of the missense predictors on the test set only (Suppl. Table 1), our tool reached the best scores in F1-Score and MCC (0.90 and 0.80, respectively) compared to other predictors with full coverage of the test set. EVE showed F1-Score and MCC values of 0.91 and 0.83, respectively, on a subset (82.5 %) of variants where it reached a prediction. By contrast, MutPred2 was able to predict every pathogenic variant as pathogenic, albeit at the cost of predicting almost half of the benign variants as pathogenic, resulting in a high number of False Positives.

Overall, Vasor outperformed other predictors consistently according to ROC and precision-recall-curves, revealing a well-balanced prediction with few False Negatives and False Positives both on the entire dataset and the test set.

### 3.4. Vasor classifies the majority of variants with high certainty

Additionally, we investigated the distribution of Vasor’s output, the probability of pathogenicity values. Vasor assigns the majority of benign cases low probability values (75% of benign variants < 0.23 probability of pathogenicity), whereas the majority of pathogenic cases are assigned a high probability value (75% of pathogenic variants > 0.76 probability of pathogenicity) (Fig. 5). Furthermore, Vasor showed no misclassifications of variants in the dataset for values below 0.24 and above 0.84, indicating high certainty for benign variant predictions in the range 0 to 0.24 (75 % of the benign variants) and pathogenic variant predictions in the range 0.84 to 1 (60 % of the pathogenic variants).

**Fig. 5:**
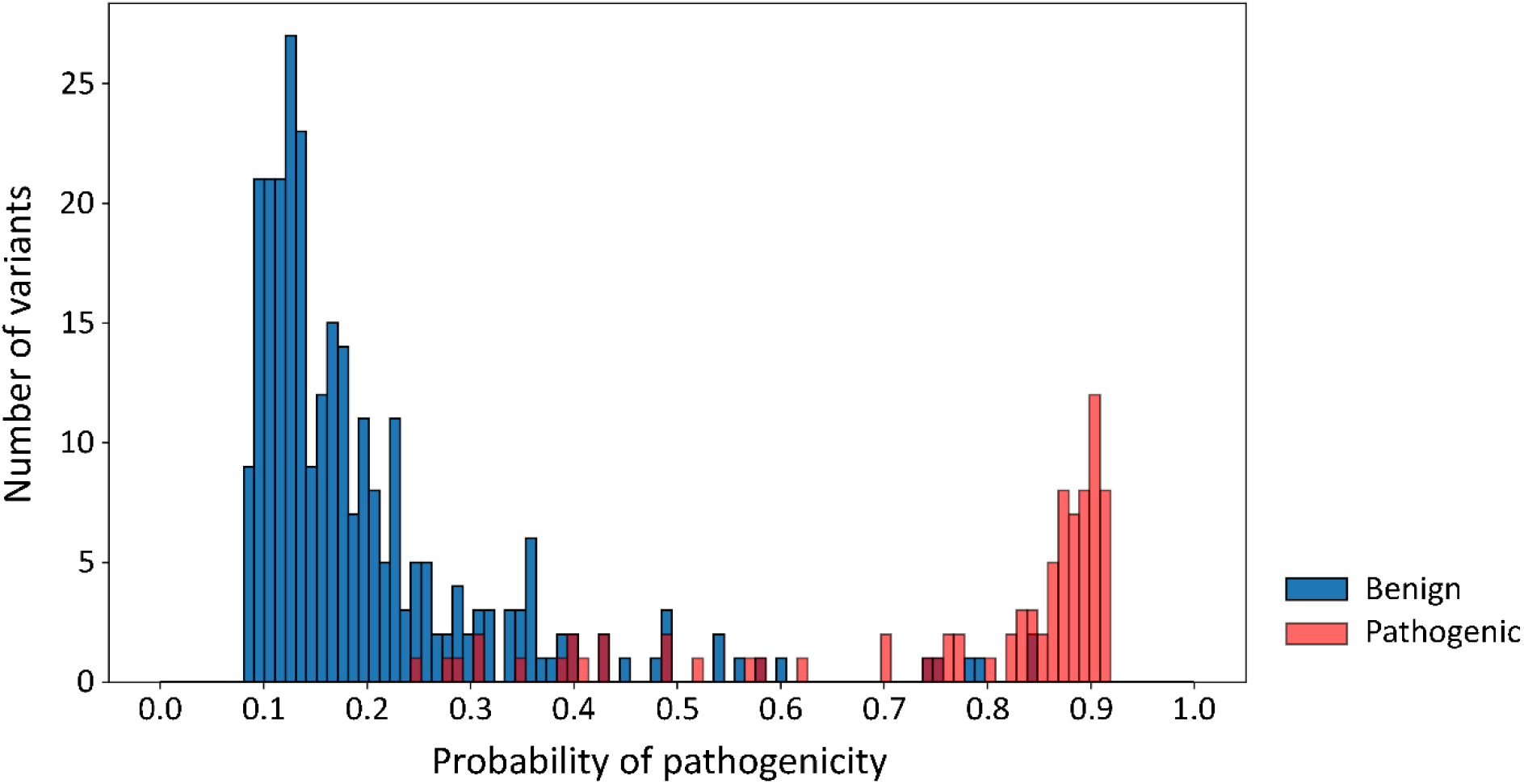
Distribution of probability of pathogenicity values over the entire dataset. Distribution of Vasor’s probability of pathogenicity output for benign (blue) and pathogenic (red) variants. Vasor classified 75 % of benign variants into the benign category with values below 0.23, which is below the lowest probability value of any pathogenic variant (0.24) within the dataset. 60 % of pathogenic variants were classified into the pathogenic category with values above 0.84, which is greater than the highest probability value of any benign variant (0.84) within the dataset. 75 % of pathogenic variants were classified with probability values greater than 0.76.

We further investigated the usage of SMOTE to generate data points for the minority class (i.e., pathogenic variants). Due to the method underlying SMOTE, SMOTE-generated data points are expected to follow the distribution of pathogenic variants within the probability of pathogenicity curve. Accordingly, no SMOTE data point was predicted with a lower value of probability of pathogenicity than 0.29, and data points mainly clustered within the high certainty zone (Suppl. Fig. 6).

Overall, Vasor showed a robust separation of probability of pathogenicity values of both variant classes, indicating that Vasor classified most variants within the dataset with high certainty.

### 3.5. Easy accessibility of Vasor as a webserver tool

Using Vasor, we precalculated the effect of every possible amino acid substitution for MDR3, resulting in a heatmap of 1286 * 20 probabilities of pathogenicity (Fig. 6, SI). We mapped the average probability of pathogenicity of each position onto the MDR3 protein structure to visualize positions that are functionally more sensitive to substitutions (Fig. 7). As expected, areas near the ATP-binding site within the nucleotide-binding domain displayed a high average probability of pathogenicity. Similarly, buried residues within the helices forming the transmembrane part showed high sensitivity as several missense mutations may lead to a disruption of the helical structure. More exposed residues located on the outsides of helices or in flexible regions, such as the small extracellular loops, displayed less sensitivity. However, this trend does not exclude that specific variants at seemingly less sensitive sites can be pathogenic, and vice versa.

**Fig. 6:**
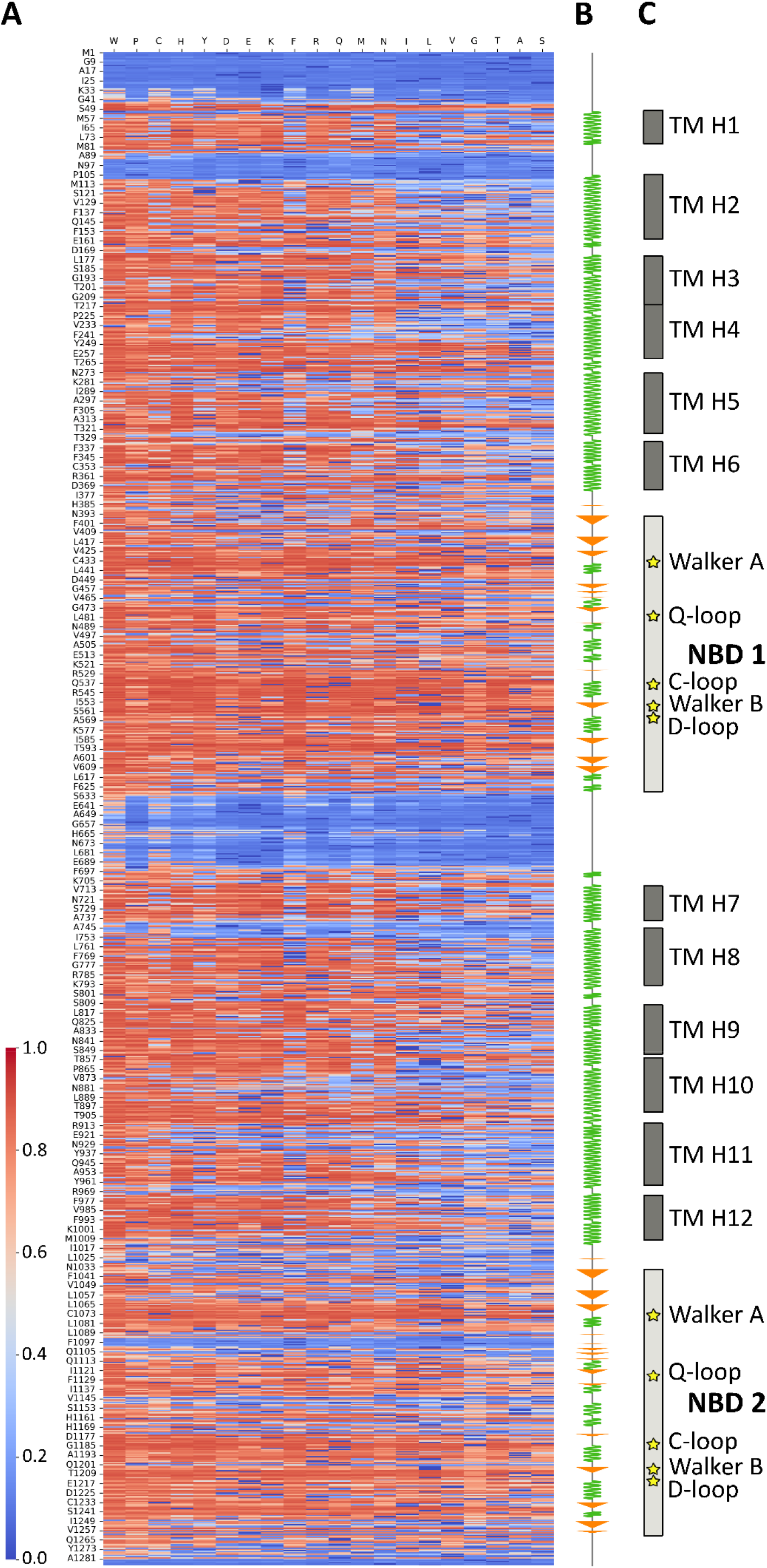
Heatmap of predictions for every possible amino acid substitution in MDR3. [A] Color-coded predictions for every position (displayed on the y-axis) within the MDR3 protein and every possible amino acid substitution (x-axis). Prediction values range from likely benign (blue) to likely pathogenic (red). [B] Secondary structure of MDR3. α-helical stretches are depicted as green zig-zag curves, β-sheet stretches as orange arrows. [C] Domains, secondary structure elements, and characteristic motives are indicated on the right. TM H: Transmembrane Helix, NBD: Nucleotide-binding domain.

**Fig. 7:**
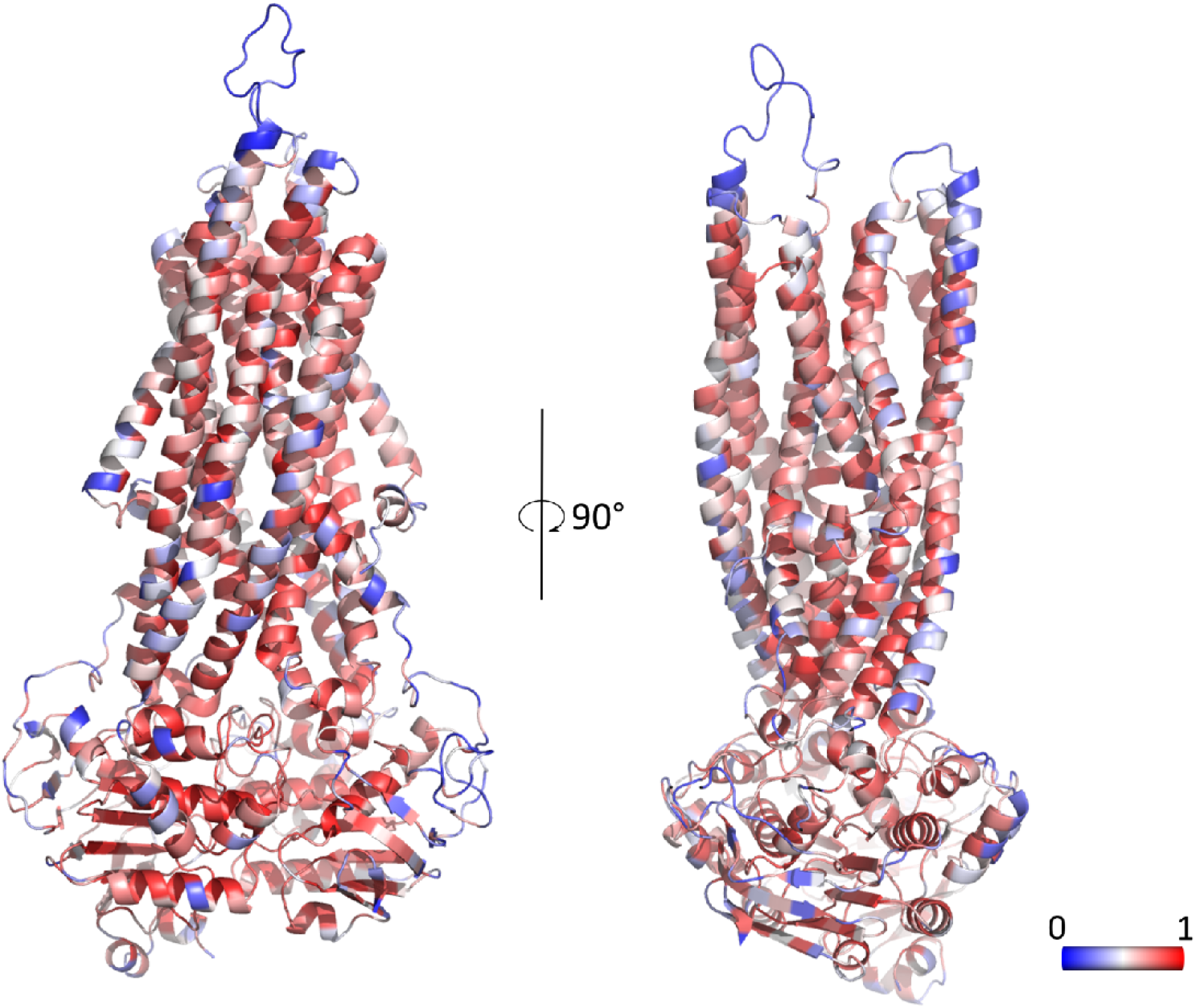
Mapping the average pathogenicity onto the structure of MDR3. Prediction values for each position were averaged over all possible substitutions. Values closer to 0 (most likely benign) correspond to blue, values closer to 1 (most likely pathogenic) correspond to red residues.

We also used the pre-computed heatmap for rapid lookup and output generation of the webserver tool, thus, eliminating waiting time for users needing a prediction for a specific MDR3 variant. The webserver can be accessed at https://cpclab.uni-duesseldorf.de/mdr3_predictor/. It requires as input an MDR3 variant (with the amino acid of the reference sequence in the one-letter format, its position within the canonical sequence of Uniprot ID P21439-1, and the substituted amino acid in the one-letter format) and yields the predicted effect of the entered variant, either benign or pathogenic, together with the probability of pathogenicity. Additionally, the variant position is depicted in the 3D structure of MDR3, and high-quality images of reference sequence amino acid, variant, and the overall MDR3 structure can be downloaded. The heatmap is also downloadable from the webserver for implementation in other applications.

## 4. Discussion

Although recent years have resulted in a plethora of general predictors for protein properties, their performance on specific proteins of interest can differ greatly (79). While existing state-of-the-art tools to predict substitution effects perform admirably on the MDR3 protein, especially EVE (17), the potential for improvement is given both for the performance on and coverage of the MDR3 dataset, since not every general predictor can classify each MDR3 variant.

To improve predictions, we created the, to our knowledge, largest dataset specific for pathogenic and benign variants of MDR3, obtained from the literature and gnomAD database and comprising 85 pathogenic and 279 benign variants. As the generation of a high-quality dataset is a critical first step for any machine learning approach (71,78), we carefully screened the literature specifically for MDR3 variants, filtering out variants with unclear disease associations. To counteract the bias that mainly pathogenic variants are chosen for detailed *in vitro* or *in vivo* analysis, we obtained variants from the gnomAD database (58). Since there may be potentially disease-associated variants in the database, we implemented an additional filtering step of removing variants categorized as likely pathogenic or pathogenic as evaluated by VarSome (59) to exclude false-negative variants. The dataset resulting from this strategy was then kept as is, i.e., no variants were added or removed, thus eliminating the potential to introduce bias from the researcher. Using established general predictors and variant site properties, we trained an MDR3-specific machine-learning model, termed Vasor, to classify protein missense variants into benign or pathogenic. Vasor outperforms general predictors: Over the entire dataset, Vasor shows F1-Score and MCC of 0.85 and 0.81, respectively, while the second-best method, EVE, follows with scores of 0.81 and 0.77, respectively, but coverage of only 85.7 %. By contrast, Vasor ensures high-quality predictions for all MDR3 missense variants. As machine learning models trained on a specific dataset exhibit a bias towards overperformance on this dataset, Vasor has an inherent advantage when evaluated on the entire dataset over other predictors. Notably, the superior performance of Vasor is also present on the independent test set, where Vasor only misclassified two (5 %) benign and two (5 %) pathogenic variants, leading to the highest performance compared to other predictors as indicated by F1-Score and MCC of 0.9 and 0.8, respectively. Although EVE and PON-P2 achieve similar performances for the test set, they only cover a fraction of the variants (82.5 % and 37.5 %, respectively). Overall, no other analyzed predictor provided a similarly good balance of consistently low False Negative and False Positive predictions. Both measures have important implications for using Vasor within a clinical setting: Predictors with a high number of False Negatives will lead to variants found within patients being falsely given no attention, whereas a high number of False Positives will result in a predictor raising too often a false alarm for an actually benign variant.

We established an easily accessible webserver for reliable and fast predictions of novel MDR3 variants based on Vasor. It can serve as an important step for deciding which variants to study and to get the first indication of a variant effect. It does not eliminate the need for classical *in vitro* studies for mutational impact, however, and in a clinical setting, the ACMG-AMP guidelines (60) should be followed. The webserver classifies single-site amino acid substitutions into the categories benign or pathogenic. Truncation, insertion, and deletion variants of MDR3 cannot be assessed. However, the probability of pathogenicity for such variants is often more definite (81,82). Of note, the effect of a single missense variant within the biological context might not always be a clear-cut pathogenic or benign effect. Therefore, the probability of pathogenicity provided by the webserver can act as an indicator of prediction reliability.

As a limitation, the exact mechanism underlying a pathogenic variant cannot be inferred from the current tool. MDR3 missense variants may impact protein folding and maturation, activity, or stability (13), and several of these categories can be influenced. Information on mechanistic dysfunction may aid in targeted therapy. In terms of machine learning, such a multi-class classification problem might be solved – with the premise of a sizeable dataset of quality-assuredvariants. Unfortunately, we are unaware of such a dataset for MDR3. The currently employed dataset strived for such quality-assured variants; however, especially lacking large-scale functional studies of benign variants, variants indicated by VarSome as of unclear significance were included. Thus, we encourage the scientific community to submit novel MDR3 variants with a proven effect on folding, maturation, activity, and stability to the authors for addition into the dataset to improve and develop Vasor further.

## 5. Acknowledgments

We are grateful for computational support and infrastructure provided by the “Zentrum für Informations- und Medientechnologie” (ZIM) at the Heinrich Heine University Düsseldorf and the computing time provided by the John von Neumann Institute for Computing (NIC) to HG on the supercomputer JUWELS at Jülich Supercomputing Centre (JSC) (user ID: HKF7, VSK33).

