## Supplemental Information for "Vasor: Accurate prediction of variant effects for amino acid substitutions in MDR3"

Address correspondence to:

**Table S1: Detailed performance measurements of Vasor in comparison to EVE, PolyPhen-2, PON-P2, and MutPred2 on the independent test set.**

| | Vasor | EVE | PolyPhen-2 | PON-P2 | MutPred2 |
|---|---|---|---|---|---|
| **Recall** | 0.90 | 0.83 | 0.95 | 0.75 | 1.00 |
| **Specificity** | 0.90 | 1.00 | 0.80 | 1.00 | 0.55 |
| **Precision** | 0.90 | 1.00 | 0.83 | 1.00 | 0.69 |
| **NPR** | 0.90 | 0.83 | 0.94 | 0.92 | 1.00 |
| **Accuracy** | 0.90 | 0.91 | 0.88 | 0.93 | 0.78 |
| **F1-Score** | 0.90 | 0.91 | 0.88 | 0.86 | 0.82 |
| **MCC** | 0.80 | 0.83 | 0.76 | 0.83 | 0.62 |
| **TP** | 18 | 15 | 19 | 3 | 20 |
| **FN** | 2 | 3 | 1 | 1 | 0 |
| **TN** | 18 | 15 | 16 | 11 | 11 |
| **FP** | 2 | 0 | 4 | 0 | 9 |
| **Coverage** | 100 | 82.5 | 100 | 37.5 | 100 |

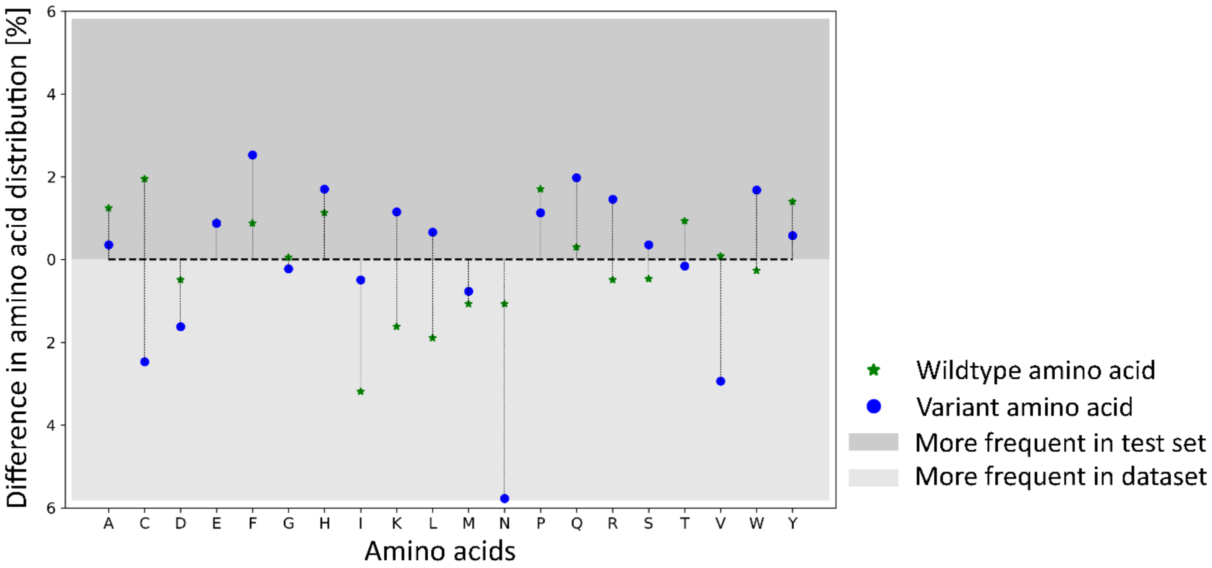


**Supplementary Fig. 1: Agreement of the distributions of amino acids between the entire dataset and the test set.** The distribution differences of reference sequence and variant amino acids between dataset and test set was computed as RMSD differences.

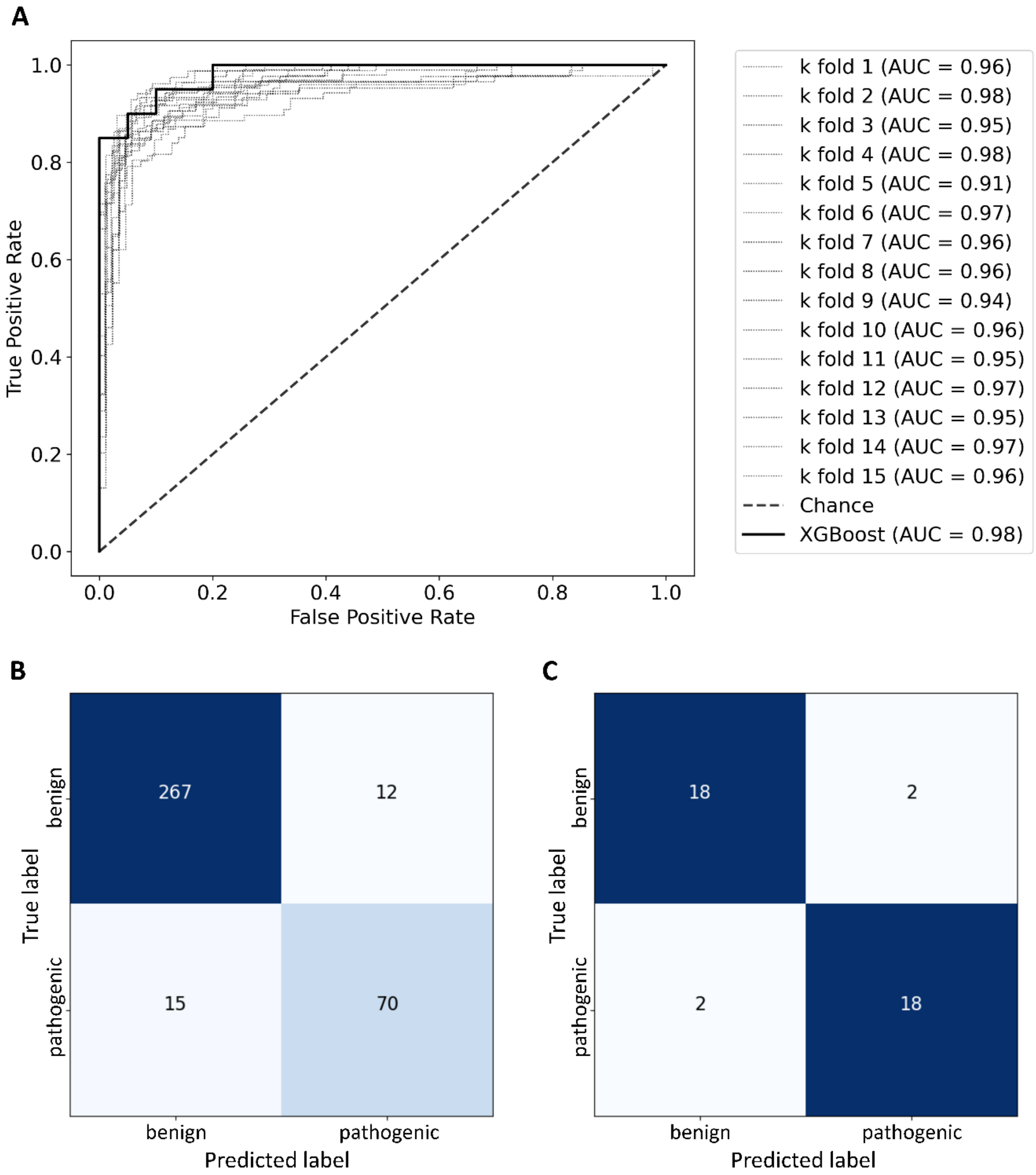


**Supplementary Fig. 2: XGBoost model performance without feature selection.** [A] ROC curve of the performance of an XGBoost model trained on every feature within the dataset. The performance of the model on the test set (solid line) is compared to the performances during the repeated *k*-fold cross-validation (dotted lines). [B] Confusion matrix of the model on the entire dataset. [C] Confusion matrix of the model on the test set.

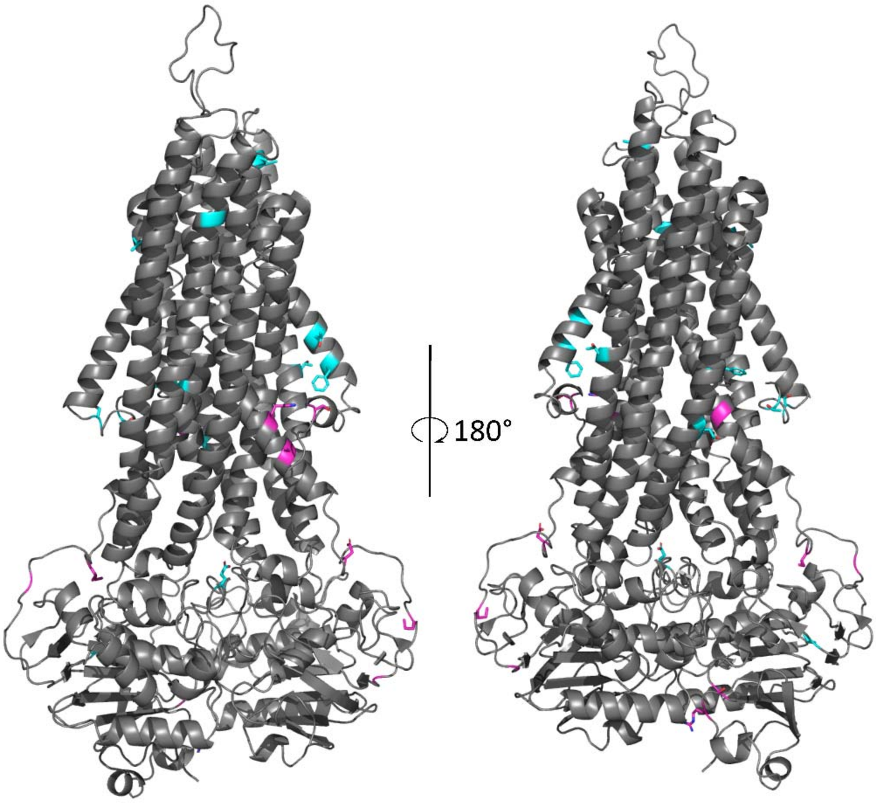


**Supplementary Fig. 3: Distribution of misclassified variants.** Misclassified variants were mapped to the MDR3 structure. Vasor-misclassified False Negatives are depicted in cyan and False Positives in pink.

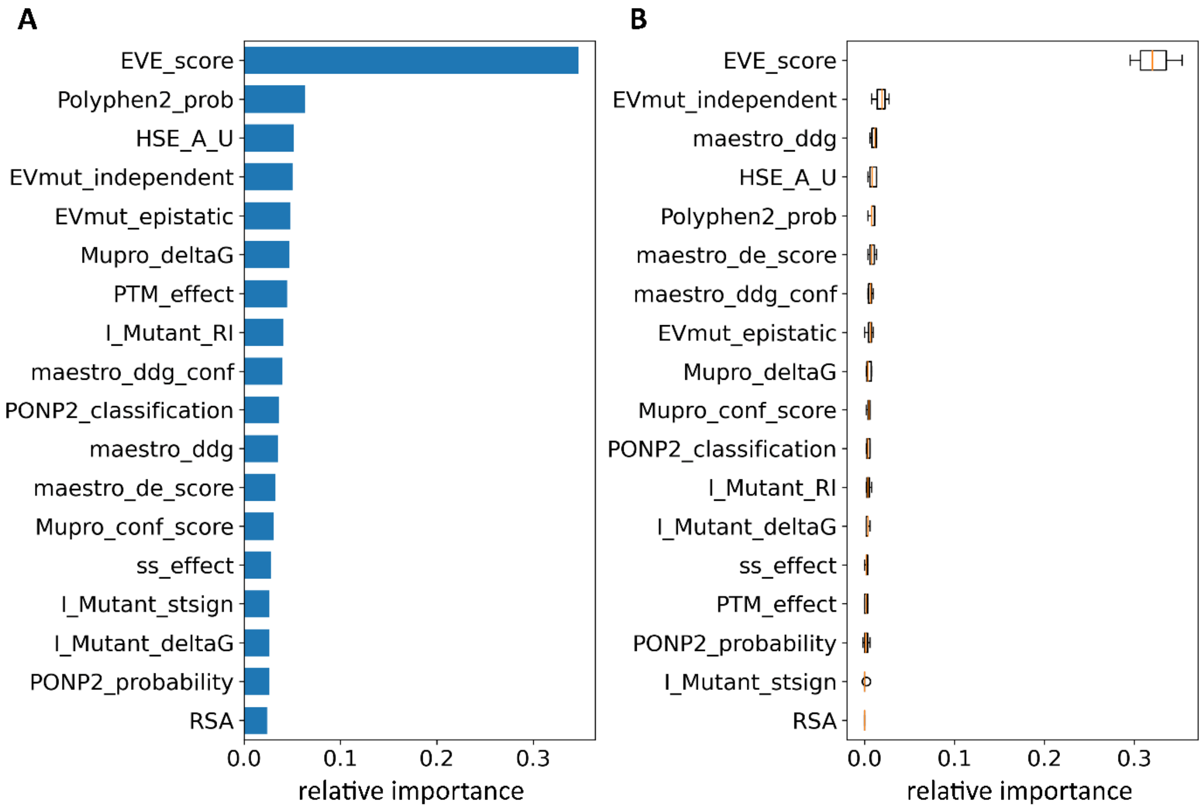


**Supplementary Fig. 4: Importance of the features.** [A] Tree-based feature importance. [B] Permutation importance. Each feature was subjected for permutation for 10 repeats. Mean values of those repeats are depicted as orange line, with the box ranging from the first to the third quartile of the data. The whiskers extend 1.5 times the inter-quartile range from the box. Outlier points located further than the whiskers are depicted as points if present.

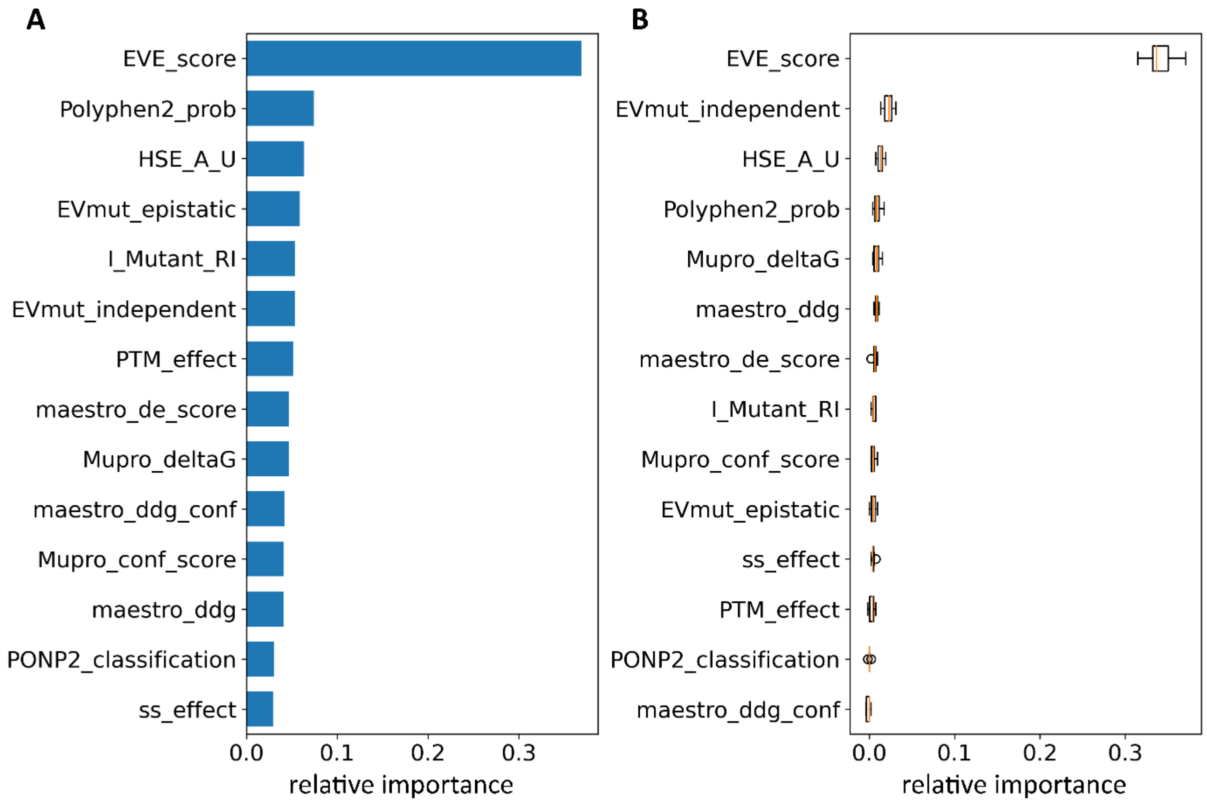


**Supplementary Fig. 5: Importance of the features in Vasor.** [A] Tree-based feature importance. [B] Permutation importance. Each feature was subjected for permutation for 10 repeats. Mean values of those repeats are depicted as orange line, with the box ranging from the first to the third quartile of the data. The whiskers extend 1.5 times the inter-quartile range from the box. Outlier points located further than the whiskers are depicted as points if present.

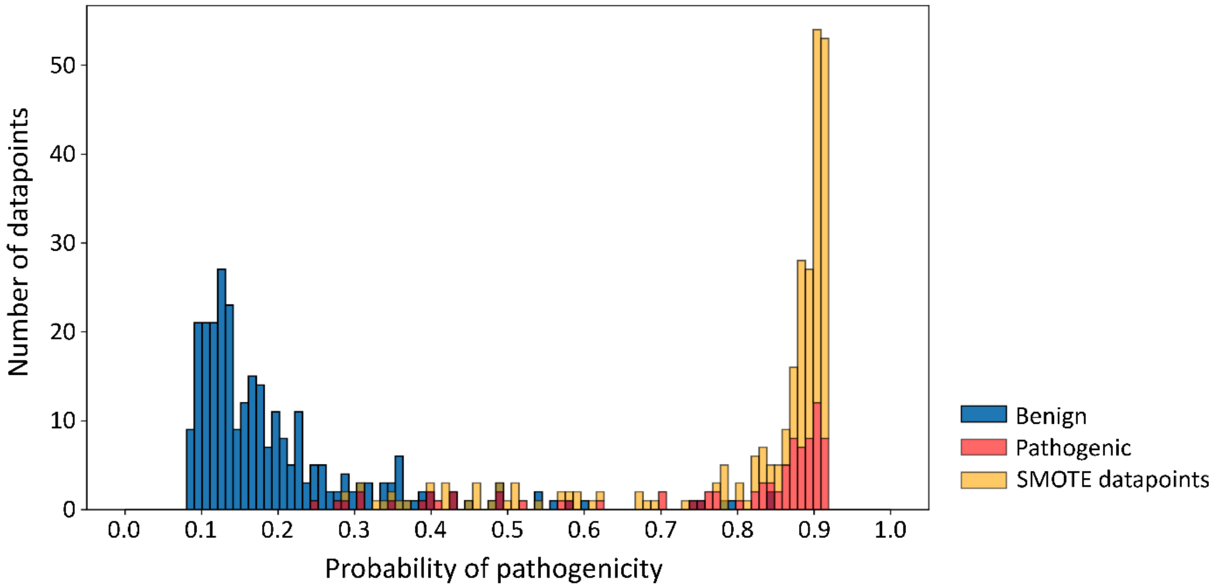


**Supplementary Fig. 6: Distribution of probability of pathogenicity values over the entire dataset including SMOTE-generated datapoints.** Distribution of Vasor's probability of pathogenicity output for benign (blue) and pathogenic (red) variants, and SMOTE-generated data points for the pathogenic class (orange). Pathogenic and SMOTE data points are represented as stacked bars. Vasor classified 75 % of benign variants into the benign category with values below 0.23, which is below the lowest probability value of any pathogenic variant (0.24) within the dataset. 70 % of pathogenic variants and SMOTE data points were classified into the pathogenic category with values above 0.84, which is greater than the highest probability value of any benign variant (0.84) within the dataset. 75 % of pathogenic variants and SMOTE data points were classified with probability values > 0.82.
